# Gene Ontology Meta Annotator for Plants (GOMAP)

**DOI:** 10.1101/809988

**Authors:** Kokulapalan Wimalanathan, Carolyn J. Lawrence-Dill

## Abstract

Annotating gene structures and functions to genome assemblies is necessary to make assembly resources useful for biological inference. Gene Ontology (GO) term assignment is the most used functional annotation system, and new methods for GO assignment have improved the quality of GO-based function predictions. The Gene Ontology Meta Annotator for Plants (GOMAP) is an optimized, high-throughput, and reproducible pipeline for genome-scale GO annotation of plants. We containerized GOMAP to increase portability and reproducibility and also optimized its performance for HPC environments. Here we report on the pipeline’s availability and performance for annotating large, repetitive plant genomes and describe how GOMAP was used to annotate multiple maize genomes as a test case. Assessment shows that GOMAP expands and improves the number of genes annotated and annotations assigned per gene as well as the quality (based on *F*_*max*_) of GO assignments in maize. GOMAP has been deployed to annotate other species including wheat, rice, barley, cotton, and soy. Instructions and access to the GOMAP Singularity container are freely available online at https://bioinformapping.com/gomap/. A list of annotated genomes and links to data is maintained at https://dill-picl.org/projects/gomap/.

## Background

Plant genomes are notably repetitive and hard to assemble. As such, long-read sequencing technologies have been quickly and widely adopted [1, 2] to enable high-quality *de novo* assembly of plant genomes. The number of plant long-read, whole-genome sequencing datasets are rapidly increasing (See table 1) and would lead to increased number of high-quality plant genome assemblies in near future. In order to make the best use of high-quality assemblies for functional genomics applications, improved computational tools for gene structure and function prediction must also be developed and adopted.

**Table 1:** Comparison of maize input sequences

| Year | Studies | Runs |
| --- | --- | --- |
| 2016 | 16 | 606 |
| 2017 | 33 | 880 |
| 2018 | 77 | 1399 |
| 2019 | 148 | 4104 |
| 2020 | 230 | 2614 |

NCBI SRA was queried on Jan 30, 2021 with the following parameters: `((("pacbio smrt"[Platform]) OR "oxford nanopore"[Platform]) AND Embryophyta[Organism] AND wgs[strategy])`

In 1998, the Gene Ontology (GO) consortium released the first common vocabulary describing gene function across species, thus enabling a genome-wide and comparative approach to functional genomics [3]. GO is divided into three categories or sub-ontologies, namely cellular component (CC), molecular function (MF) and biological process (BP). Various tools and approaches were developed to assign GO terms to genes, and a raft of statistical methods to interpret high-throughput experimental results for GO-based gene function implications were developed and released [4–8]. More recently, the Critical Assessment of protein Function Annotation (CAFA) competition has enticed research groups to develop tools that improve the accuracy and coverage of gene function prediction [5, 7, 9]. Unfortunately, methodologies developed through CAFA have not been widely adopted for annotating plant genomes, and existing plant-specific GO annotation pipelines mainly focus on subsets of GO terms rather than the full set of terms available [10, 11].

We sought to assess the performance of some of the best-performing methods produced through CAFA1 for assigning gene function to plant genomes and to produce an improved functional annotation dataset for maize. These efforts were successful, with improvements to prediction outcomes measured in terms of precision, recall, and coverage [12]. Obvious next steps were to generalize the developed pipeline and to apply it to other maize lines and additional plant species then to evaluate its performance for annotating gene function to those genomes. Here we present GOMAP (Gene Ontology Meta Annotator for Plants) pipeline that generalizes the methods used to produce the maize-GAMER datasets, with improvements to computational performance, reproducibility, and portability. We also present the utility of GOMAP by annotating genomes assemblies of four maize inbred lines, namely B73 RefGen v4 (B73v4), W22, PH207 and Mo17 [1, 2, 13, 14]. The GOMAP annotations are compared to the community annotations for B73v4 and PH207. Gramene annotated B73v4 RefGen v4 using Ensembl Compara method and produced a high-confidence and high-coverage GO annotations [15].

The PH207 dataset was obtained from the supplemental tables of the PH207 genome sequencing paper [13]. GO terms for PH207 genes were annotated using InterProScan v5.0 that uses domain presence to assign GO terms to input protein sequences[13]. We compared GOMAP to community methods used to annotate B73v4 and the PH207 to illustrate the differences among datasets produced by three approaches for GO annotation.

## Materials and Methods

### Overview of the Annotation of input sequences

The GOMAP uses sequence-similarity, domain-presence and mixed-method pipelines to annotate GO terms to the input protein sequences to produce a single unique and non-redundant GOMAP aggregate dataset as the result (See Figure 1). Sequence similarity searches are performed against two plant datasets: Arabidopsis and UniProt. The Arabidopsis dataset contains protein sequences downloaded from TAIR and curated GO annotations [16]. The UniProt dataset contains protein sequences from the top plants species that were ranked by number of curated GO annotations available in UniProt [17]. The first set of annotations is generated using BLAST to obtain reciprocal-best-hits between input and Arabidopsis sequences, and inheriting curated GO terms from Arabidopsis to the input sequence [18]. A second set of annotations is obtained using a similar approach, but instead of Arabidopsis the search is performed against the top ten annotated plant species in the UniProt database. Presence of valid domains in the input sequences is identified using the InterProScan5 pipeline. InterProScan uses fourteen types of protein signatures to detect putative domains in the input sequences, and assign GO terms [19]. As per documentation, InterProScan only reports valid domains and GO annotations for the valid domains, so the GO annotations are not filtered based on scores for this step. Three mixed-method pipelines from the first iteration of the CAFA competition (CAFA1 tools) are used to annotate GO terms to the input sequences, namely Argot2.5, FANN-GO and PANNZER [7, 20–22]. Two CAFA1 tools require preprocessing of input sequences before they can be used to annotate GO terms. Argot2 requires the BLAST hits of the input sequences to the UniProt database and Pfam hits identified by HMMER search against the Pfam domain database [23–25]. PANNZER only requires the BLAST hits to the UniProt database for the annotation process. The 6 annotation datasets generated from previous steps are aggregated. Next any redundancy or duplication introduced by aggregation is removed to produce a final aggregate dataset. See Defoin-Platel et al. for the definitions of redundancy and duplication, and maize-GAMER for more details about the annotation methods used in GOMAP [12, 26]. For the analyses described here, non-plant-specific annotations were not removed. See the accompanying GitHub repository for a R script that can be used to filter for plant specific terms. The removal of non plant-specific GO terms did slightly reduce the number of annotations per GO category (See Supplentary table ST2). The GOMAP and community annotations retained about 99% the original annotations. This enables researchers to use such terms to formulate novel hypotheses about potential plant gene functions that could be inspired by data obtained in non-plant systems (e.g., genes involved in the initiation of neurons could be involved in initiation of root hairs, information on flagellar function in lung cells could inform ideas on flagellated sperm function in gymnosperms, etc.).

**Figure 1:**
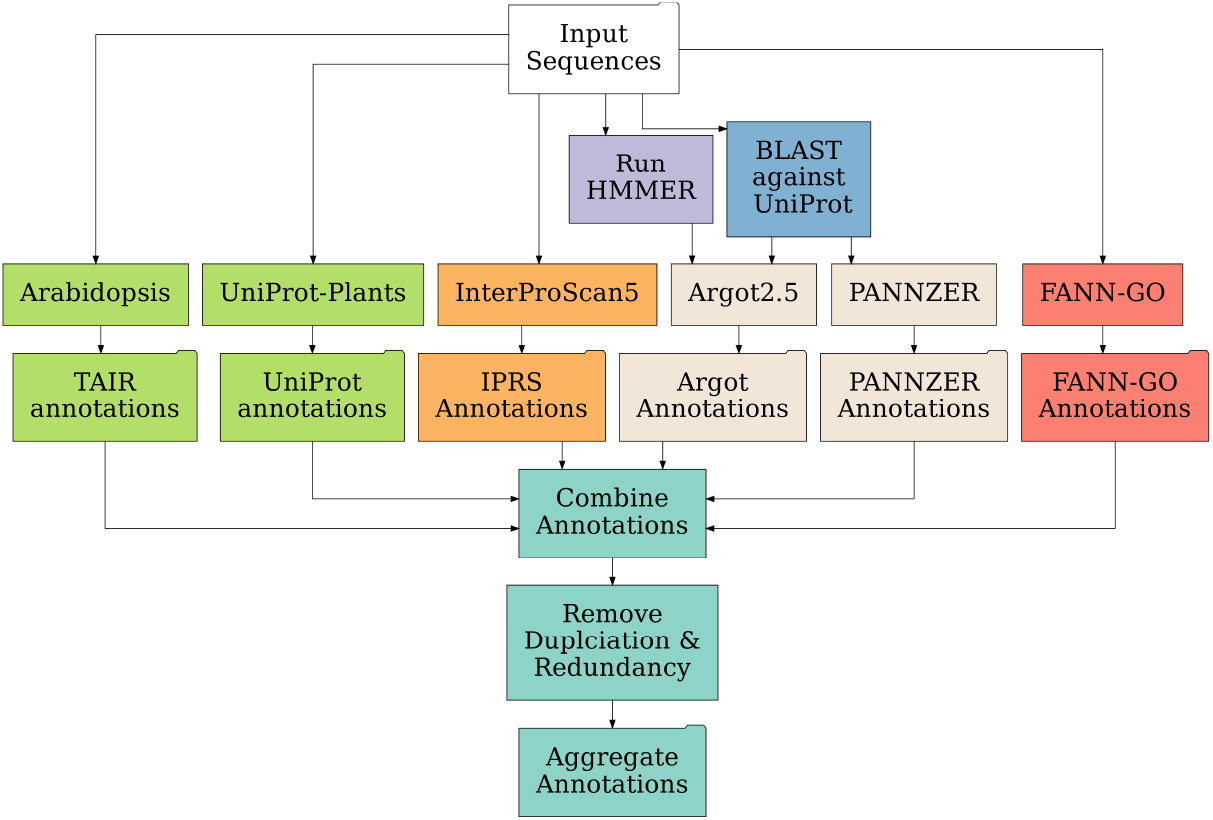
Overview of processes used to create GOMAP Annotations. Top: data inputs are shown as a white box. Sequence similarity components are shown in lime green. InterProScan domain-based annotations are shown in orange. For the CAFA processes (Argot2.5 and PANNZER; shown in tan boxes) pre-processing steps are shown in purple and blue, respectively. The CAFA process FANN-GO does not require preprocessing (red). Once each annotation type is produced, these are combined, duplicates and redundancies are removed, and the aggregate dataset is assembled (turquoise).

**Figure 2:**
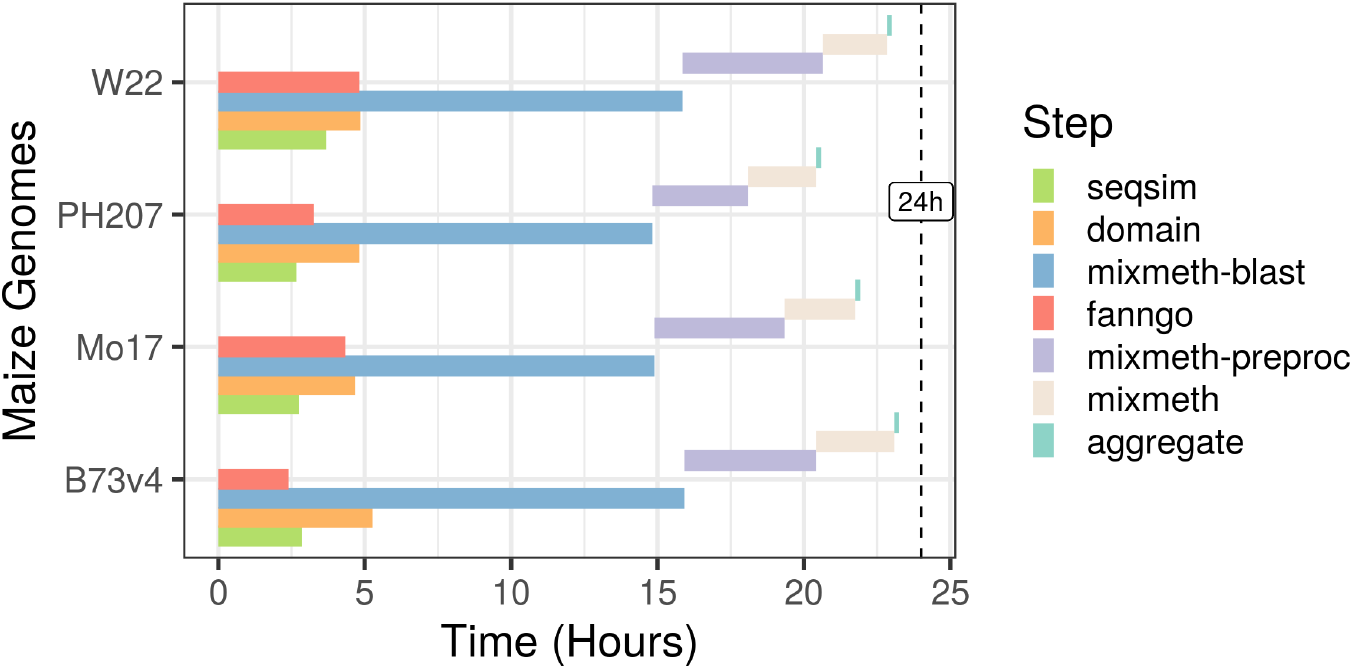
Comparison of runtime for GOMAP steps across four maize genomes. Steps are color-coded as shown in the figure key at right. Start time occurs at zero. The four steps shown simultaneously within a single maize genome (i.e., seqsim, domain, mixmeth-blast, and fanngo) run in parallel. For all maize genomes, the full annotation time took less than 24 hours on the PSC Bridges cluster.

### Implementation and Containerization of GOMAP

The GOMAP pipeline was developed by containerizing the refactoring maize-GAMER code into one singularity container [12, 27]. The GOMAP Pipeline is implemented in Python and R. Python code is used to run open-source tools for annotating GO terms, and R code is used to aggregate and clean annotation results. GOMAP was containerized to improve usability, portability and reproducibility. Containerization eliminates the need to install and configure dependencies. Singularity containerization was chosen because it works seamlessly in high performance computing (HPC) environments [27], and it has been widely adopted by HPC clusters. Several bottlenecks were encountered when containerizing GOMAP: the large size of the pipeline, long runtime on a single machine, and the use of MySQL and MATLAB by mixed-method pipelines.

The uncompressed data required for GOMAP pipeline uses about 110 GB of local disk space. This large size is due to the inclusion of external tools and data, which results in a large container that creates issues during the development and distribution of GOMAP via free public resources. Some tools such as PANNZER were dependent on a back-end MySQL database, and FANN-GO included MATLAB specific code for the annotation. These two components complicated the complete containerization and subsequent efforts to run GOMAP on HPC systems. The original PANNZER code was updated to use a SQLite database, and the SQLite database file works seamlessly in HPC systems eliminating the need for MySQL. The FANN-GO code was updated to use open source GNU Octave instead of MATLAB. The ability to include Octave in the container enabled GOMAP to be run on any HPC system and completely enclose all the data and software required for GOMAP in the container. The pre-built GOMAP containers are currently shared via CyVerse^[1]^ and GitHub^[2]^ [28]. Run time for GOMAP on a single machine on a single node in the Iowa State University HPC Condo Cluster^[3]^ for 40,000 protein sequences takes between 12-14 days. To improve runtime, GOMAP was separated into different steps that run concurrently. Moreover, the 2 steps with longest runtime, InterProScan (1-2 days) and BLAST search against the UniProt sequence database (8-10 days), were parallelized. Most HPC environments have shorter time limits (e.g. 2-5 days), so parallelizing is necessary to complete these steps within such limitations.

### Annotation of maize genomes as a test case

Two versions of the Maize B73 reference genome releases were annotated for the maize-GAMER project. At the completion of containerizing GOMAP, genomes of three more maize inbred lines had been released: W22, Mo17, and PH207. The GOMAP container was used to annotate the three newly released genomes and replicate the annotations for B73 RefGen v4. The input files were downloaded from MaizeGDB as shown in Table **ST1**. The protein sequences downloaded for each genome were filtered to retain only the longest translated transcript for each gene. Each input fasta file from the 4 different maize genomes was annotated by GOMAP on the Pittsburgh Supercomputing Center (PSC) Bridges HPC Cluster ^[4]^. GOMAP steps were run on Regular Shared Memory nodes. Each regular shared memory node is configured with two Intel Haswell (E5-2695 v3) CPUs (28 Total CPU cores) and 128 GB memory. In addition, two community annotation datasets for B73 RefGen v4 and PH207 were obtained for comparison. The community annotation for B73 RefGen v4 was downloaded from Gramene using GrameneMart tool [15, 29]. The community annotations for PH207 (PH207-community) were obtained from the supplemental methods of the original publication by Hirsch et al. [13]. The GOMAP datasets for the four inbred lines and the two community datasets were used for downstream comparison and evaluation.

### Assessment and comparison of analysis and evaluation metrics for maize annotation datasets

Maize annotation datasets were assessed using two different metrics: analysis metrics and evaluation metrics. Analysis metrics were used to assess and compare the quantity of the annotations among the datasets, whereas the evaluation metrics assess the quality of the annotations by comparing against a gold-standard dataset produced by manual curation. The data and R code used to evaluate the datasets are available via GitHub ^[5]^.

Three analysis metrics, coverage, number of annotations normalized by gene count (i.e., number of annotations), and, were used for the assessment and comparison of maize annotation datasets. Coverage represents the proportion of the total genes that have at least one GO annotation in the predicted dataset. The number of annotations represent the total number of annotations divided by the number of genes with at least one GO annotation. The specificity for a single annotation is calculated by counting the number of ancestral terms, and the mean specificity for all annotations represents the specificity of a dataset. See Defoin-Platel et al. for detailed definitions of the analysis metrics [26]. A general comparison of analysis metrics were performed for the four GOMAP and two community maize annotation datasets. As the next step, annotations were separated into each GO category and analysis metrics were calculated and compared for each GO category. The same GO-category-specific approach was used for the generation and comparison of evaluation metrics.

A set of gold-standard annotations are required to calculate evaluation metrics. The gold-standard dataset used in maize-GAMER that was obtained from MaizeGDB was curated for the B73 RefGen v3 gene models and not for B73 RefGen v4 nor other inbred lines [30]. However, MaizeGDB has assigned gene models from B73 RefGen v3 to other inbred lines’ gene models and created a cross reference file^[6]^. This cross reference file was used to inherit curated GO terms from B73 RefGen v3 to other inbred lines and create gold-standard datasets for all four inbred lines used in this project. The R script that was used to assign the GO terms is available as part of the GitHub repository. The gold-standard GO terms inherited from B73 RefGen v3 to B73 RefGen v4, Mo17, PH207, and Mo17 were used to calculate the protein-centric evaluation metrics defined by Clark and Radivojac [31] and used for the CAFA [5, 7, 9, 32]. The three protein-centric evaluation metrics calculated were Precision (*Pr*), Recall (*Rc*) and *F*_*max*_.

### Comparison of the GOMAP and the Community, and gold-standard annotations

The comparison of maize annotations produced by GOMAP to the community annotations was restricted to the two inbred lines that had community annotation datasets, namely B73v4 and PH207 [1, 13]. Analysis and evaluation metrics were generated for both datasets and compared to GOMAP-derived datasets. In addition, gold-standard annotations were overlapped with predicted annotations from the community and GOMAP datasets and directly compared. The gold-standard terms that contained only leaf terms were expanded to include all the ancestral terms to the root node, and the same expansion was performed for the predicted annotations. The intersection of gold-standard and predicted terms was performed in three types of objects: gold-standard genes, gold-standard GO terms, and gold-standard annotations. This analysis was used to identify the gold-standard genes, GO terms and GO annotations that were found in both predicted datasets (GOMAP and Community), only in one predicted dataset (GOMAP or Community), or not found in either dataset (only gold-standard). This comparison was performed separately for each GO category for both B73 and PH207.

## Results

### Annotation of Maize Genomes using GOMAP

The GOMAP container was tested by annotating GO terms to the protein coding genes of four maize inbred lines (B73, Mo17, W22, and PH207). The size the the number protein coding genes were similar among the maize lines as expected (See Table 2). The total predicted protein coding length varied slightly among the inbred lines. W22 has the highest total length, and B73v4 has the longest. The shortest genes that were annotated are less than five amino acids long in all inbreds except B73v3. These are potential annotation errors in the database, but are reported as valid gene models. The median and mean length of the genes in the annotations are similar but vary within a narrow range, and PH207 has the lowest median and mean gene length. Three inbred-lines have longest genes that are over 5000 amino acids long. The genes that are smaller than 50 amino acids present a challenge to predicting GO terms. Mo17 had the highest proportion of genes smaller than 50 amino acids in length (*>*1300), which incidentally has the lowest annotated gene count. All the other inbred lines have less than 1% of genes shorter than 50 amino acids.

**Table 2:** Comparison of maize input sequences

| Inbred | Gene Count <sup>1</sup> | Total AA | Length |  |  |  | Small Proteins(%)<sup>2</sup> |
| --- | --- | --- | --- | --- | --- | --- | --- |
|  |  |  | min | mean | median | max |  |
| B73v3 | 39,475 | 14,382,005 | 25 | 364.33 | 306 | 4,743 | 0.56 |
| B73v4 | 39,324 | 15,373,604 | 2 | 390.95 | 316 | 5,267 | 0.86 |
| Mo17 | 38,620 | 14,640,283 | 5 | 379.09 | 306 | 5,426 | 3.44 |
| PH207 | 40,557 | 14,311,872 | 2 | 352.88 | 280 | 4,947 | 0.33 |
| W22 | 40,690 | 15,439,503 | 1 | 379.44 | 304 | 5,426 | 0.25 |

<sup>1</sup>This indicates the protein coding genes
<sup>2</sup>Any protein smaller than 50 amino acids was classified as small

### Run times of GOMAP steps for different maize genomes

Run times for GOMAP were determined using the PSC Bridges HPC cluster. The manual annotation process of maize-GAMER is complex with over 40 interdependent steps required for end-to-end annotation of a plant genome. To make the annotation process intuitive and convenient, the GOMAP annotation process combined the maize-GAMER steps into just seven discrete steps (see Table 3). The first four steps, seqsim, fanngo, domain, and mixmeth-blast, are setup to be run concurrently as independent processes. The last three steps, mixmeth-preproc, mixmeth, and aggregate, depend on the output of the first four steps. The total time taken to complete the annotation of the maize genomes were between thirty-three and thirty-six hours. The total predicted protein length and gene number had negligible impact on the total runtime of GOMAP for maize genomes, though runtimes of steps were impacted by the load of the cluster. Two parallelized steps, domain and mixmeth-blast, ran longer than other steps, but the runtime has been considerably shortened compared to the un-parallelized versions. The domain step runs for over five days without parallelization and mixmeth-blast runs for over ten days without parallelization. Notably, running steps 1-4 concurrently allows GOMAP to complete the annotation of maize genomes within twenty-four hours for each genome tested.

**Table 3:**
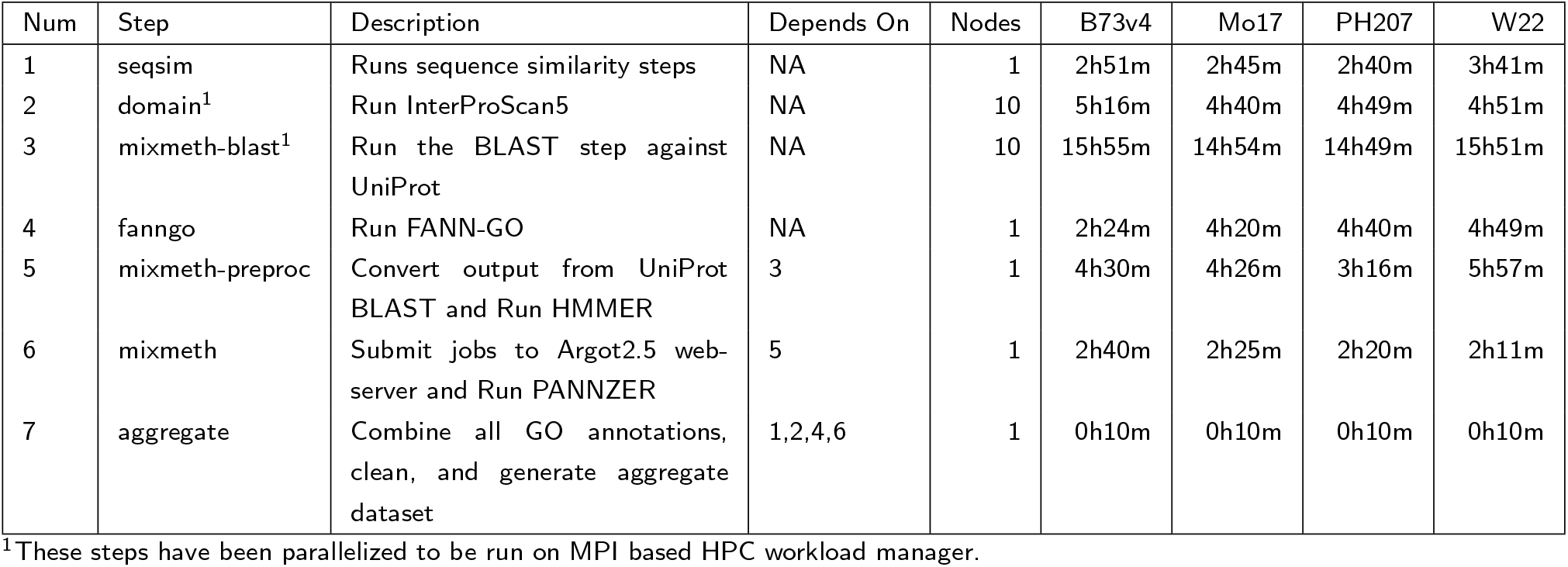
Comparison of the runtime of different GOMAP steps on PSC Bridges Cluster

### Assessment and comparison of the analysis metrics for maize annotations

Coverage, number of annotations, and specificity (See Table 4) were calculated for the GOMAP and community datasets. High coverage of around 100% is observed for all GOMAP datasets. In comparison, the community datasets for B73v4 and PH207 have about 77% and 45% overall coverage, respectively. The gold-standard datasets only cover around 3-4% of genes and provide only a smaller number of genes to calculate the CAFA evaluation metrics (Table 4). The annotations were separated by each category to get a more clear picture of the coverage (See Figure 3). The coverage changes substantially across the categories for all datasets. The GOMAP datasets have the highest coverage in the biological process category for all inbred lines (i.e., ∼100%), and have lower coverage other categories (CC:86-92%; MF:82-95%). However, both community datasets have highest coverage in the molecular function category. The PH207 community dataset had the lowest coverage among annotation datasets in all three GO categories, and the PH207-community dataset covered only about ∼10% genes in the cellular component category. The Gramene dataset had higher coverage than the PH207-community, but had lower coverage than GOMAP in all GO categories. This indicates that GOMAP produces higher-coverage datasets than both Gramene or PH207-community methods.

**Table 4:** Analysis metrics of GOMAP annotations for maize genomes.

| Inbred | Source | Total Genes | Coverage (%) |  | Annotations/Gene |  | Specificity |  |
| --- | --- | --- | --- | --- | --- | --- | --- | --- |
|  |  |  | Curated | Predicted | Curated | Predicted | Curated | Predicted |
| B73v4 | GOMAP | 39,324 | 1.26 | 11.46 | 3.54 | 100.00 | 11.99 | 10.85 |
| B73v4 | Community | 39,324 | 1.26 | 3.95 | 3.54 | 76.94 | 11.99 | 12.80 |
| Mo17 | GOMAP | 38,620 | 1.26 | 11.53 | 3.42 | 100.00 | 11.99 | 10.85 |
| PH207 | GOMAP | 40,557 | 1.26 | 11.48 | 3.34 | 100.00 | 12.05 | 10.67 |
| PH207 | Community | 40,557 | 1.26 | 2.21 | 3.34 | 45.44 | 12.05 | 11.30 |
| W22 | GOMAP | 40,690 | 1.24 | 11.57 | 3.08 | 100.00 | 12.09 | 10.78 |

**Figure 3:**
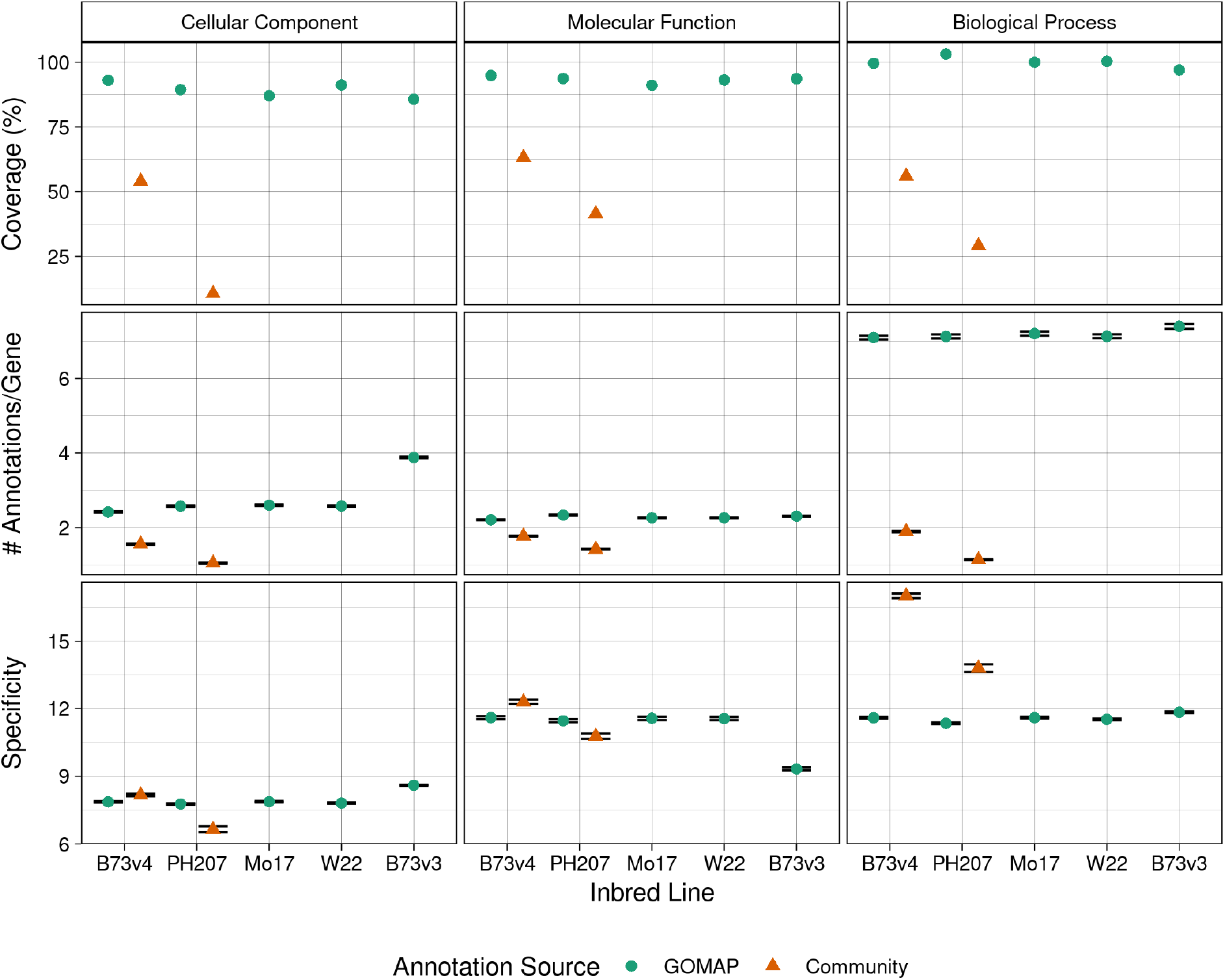
Analysis metrics calculated for the maize annotations from community and GOMAP annotations. Left column: Cellular Component. Middle column: Molecular Function. Right column: Biological Process. Top row: Percentage of genes with an annotation. Middle row: number of annotations per gene. Bottom row: specificity of the annotations. Inbred lines are denoted along the x-axis. GOMAP annotations are denoted by a green circle. Community annotations are denoted by a orange triangle. Coverage is shown as an overall percentage, but both number of annotation per gene and specificity are represented as mean values across all annotations in the dataset. Error bars indicate standard error. The confidence interval is very small so the high and low error bars overlap each other for most datasets.

The number of annotations were normalized by dividing the total number of annotations by the number of genes. This normalization allows for comparison among different datasets for the same genome and different genomes. The number of annotations vary among the inbred lines and datasets. B73v3 has the highest number of annotations among the GOMAP datasets, even though W22 had the highest number of protein-coding genes. GOMAP datasets had the highest number of annotations across all inbred lines, followed by the community datasets. The gold-standard datasets had the lowest number of annotations by a significant margin (See Table 4). In some inbreds, such as B73v4 and Mo17, GOMAP had nine times the number of annotations than the corresponding gold-standard dataset. The community datasets also have higher number of annotations than the gold-standard datasets, but the magnitude of difference was lower (∼1-3x). The number of annotations were separated by GO category and compared among each other. This allowed for the number of annotations to be compared among different inbred lines, annotation sources, and GO categories. The highest number of annotations was seen in GOMAP datasets in the BP category (∼7 annotations per gene), which is significantly higher than the community datasets in BP (B73v4:∼3x; PH207∼6x) and GOMAP datasets in other GO categories. GOMAP datasets have a higher number of annotations compared to the community datasets in all GO categories, but the magnitude of difference is not as high in CC and MF categories. The PH207 community dataset shows the lowest number of annotations across all three GO categories, and this number is especially low in the CC category. In comparison, GOMAP shows lowest number of annotations in the MF category. Gramene datasets for B73v4 has the highest number of annotations in MF and has the lowest in CC.

Specificity indicates the number of ancestral terms for a given annotations given the GO hierarchy, and the mean of all annotations for a particular dataset. Specificity represents a measure of information provided by a specific term. This metric is higher in the community datasets and gold-standard datasets in all three categories (see Table 4), compared to coverage and number of annotations. The Gramene dataset for B73v4 has higher specificity than even the gold-standard dataset. The GOMAP datasets also had lower specificity than gold-standard datasets. A more detailed analysis separated by each GO category allowed similar comparisons for coverage and number of annotations (see Figure 3). All datasets had higher specificity in BP and MF categories than CC. The Gramene B73v4 dataset has highest specificity across all GO categories, but achieved significantly higher specificity in the BP category. The PH207 community dataset has higher specificity than GOMAP only in BP category, but GOMAP has slightly higher coverage in both CC and MF categories.

### Assessment and comparison of the evaluation metrics for maize annotations

The evaluation metrics were calculated by comparing the predicted annotations to the gold-standard datasets. Three protein-centric evaluation metrics from CAFA were used to assess the annotations: Precision (*Pr*), Recall(*Rc*), and *F*_*max*_. Precision measures the proportion of predicted annotations that overlap gold-standard. Recall measures the proportion of gold-standard annotations that are correctly predicted. *F*_*max*_ is the harmonic mean of *Pr* and *Rc* and provides a single number for comparison among different methods. The evaluation metrics were calculated separately for each GO category (see Figure 4). An important factor to notice is the total number of gold-standard annotations are imbalanced and are skewed toward the CC category (See Table 5). This skewed distribution of gold-standard data directly affects the calculation of the evaluation metrics, and this is indicated by the wider standard error bars seen in MF and BP categories in Figure 4. Evaluation metrics compare the performance of the methods used for annotation, thus the following conventions are used to describe the annotation methods for maize datasets. The community method used to annotate B73v4 is called “Gramene” and the community method used to annotate PH207 is called “PH207-community” in the following section.

**Table 5:** Number of gold-standard genes, GO terms, and annotations that were assigned by GOMAP, the community annotation, and the gold-standard.

| Inbred Line | Type | Cellular Component |  |  | Molecular Function |  |  | Biological Process |  |  |
| --- | --- | --- | --- | --- | --- | --- | --- | --- | --- | --- |
|  |  | Genes | Terms | Annotations | Genes | GO Terms | Annotations | Genes | Terms | Annotations |
| B73v4 | Both | 980 | 64 | 8,014 | 56 | 183 | 518 | 126 | 271 | 1,305 |
|  | GOMAP<sup>a</sup> | 317 | 4 | 5,013 | 2 | 23 | 31 | 15 | 85 | 504 |
|  | Community<sup>b</sup> | 2 | 0 | 94 | 0 | 1 | 36 | 0 | 22 | 35 |
|  | Curated<sup>c</sup> | 14 | 34 | 2,703 | 0 | 32 | 57 | 1 | 206 | 1,184 |
|  | Total | 1313 | 102 | 15824 | 58 | 239 | 642 | 142 | 584 | 3028 |
| PH207 | Both | 151 | 22 | 784 | 40 | 100 | 283 | 71 | 77 | 304 |
|  | GOMAP<sup>a</sup> | 1,104 | 46 | 11,712 | 15 | 94 | 245 | 63 | 260 | 1,319 |
|  | Curated<sup>c</sup> | 24 | 34 | 2,987 | 0 | 41 | 96 | 1 | 234 | 1,194 |
|  | Total | 1279 | 102 | 15483 | 55 | 235 | 624 | 135 | 571 | 2817 |

<sup>a</sup>The gold-standard data overlaps with only GOMAP<sup>b</sup>The gold-standard data overlaps with only Community<sup>c</sup>The gold-standard data does not overlap with GOMAP or Community

**Figure 4:**
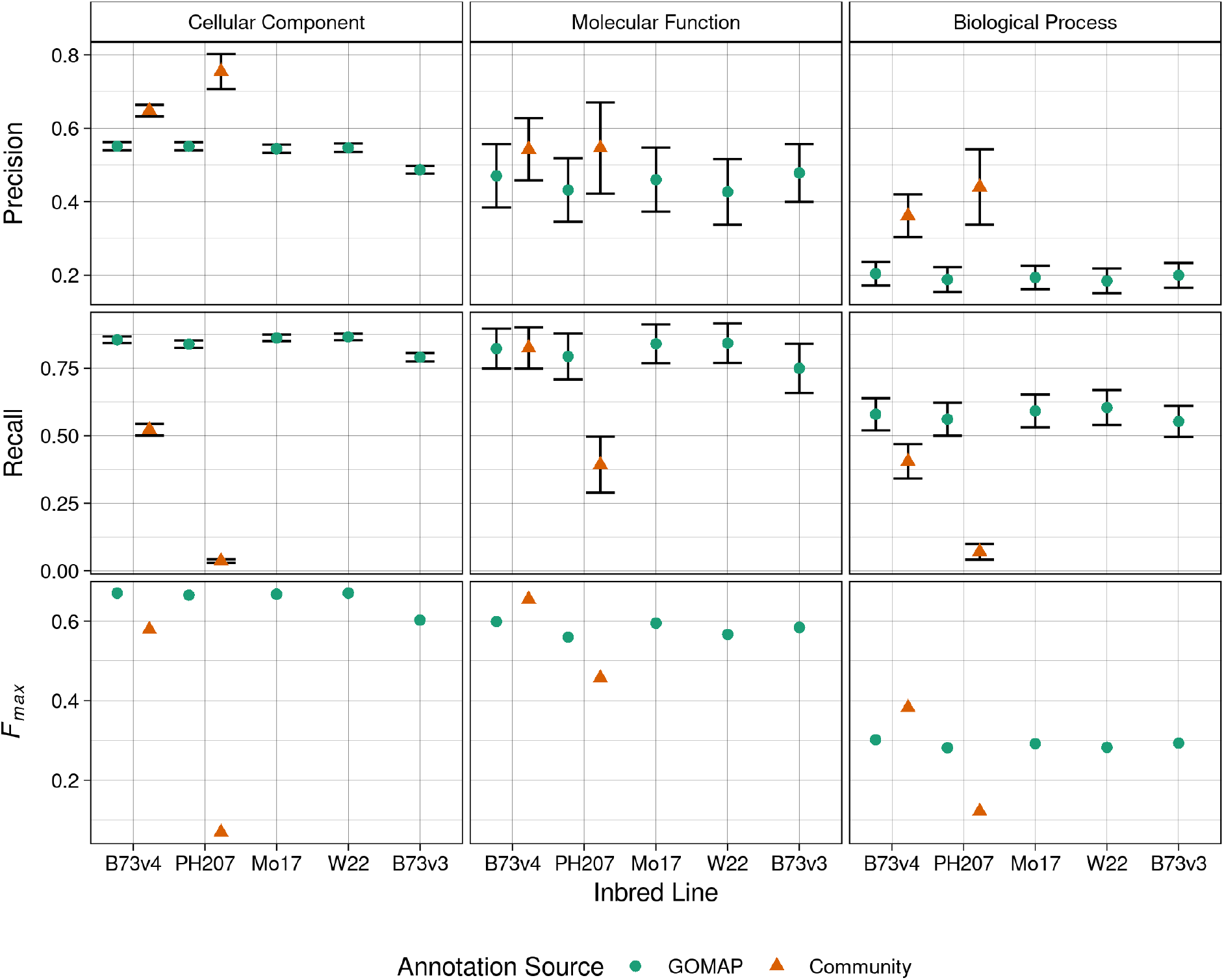
Evaluation metrics calculated for the maize annotations from the community and GOMAP. Left column: Cellular Component. Middle column: Molecular Function. Right column: Biological Process. Top row: Precision. Middle row: Recall. Bottom row: *F*_*m*_*ax*. Inbred lines are denoted along the x-axis. GOMAP annotations are denoted by a green circle. Community annotations are denoted by a orange triangle. Precision and recall are shown as the mean value of all annotations with error bars indicating standard error, but *F*_*m*_*ax* is represented as an absolute value for a specific dataset.

All methods had higher Precision in the CC category compared to other categories, while BP category had the lowest Precision overall. PH207-community method achieved the highest Precision among all datasets in all three GO categories. Furthermore, PH207-community has substantially higher Precision than GOMAP in CC and BP categories. Gramene also obtained higher precision for B73v4 than GOMAP in all three categories, although the magnitude of difference was lower. The method employed by the PH207-community is more precise in comparison to other methods. Recall values did not show a clear performance trend as seen with Precision. The recall performance varied among the methods, and no single method performed better than other methods across all GO categories. GOMAP achieved better Recall in both CC and BP categories, but Gramene showed slightly better recall (GOMAP=0.8229433; Gramene=0.8250246) than GOMAP in the MF category. It was clear both GOMAP and Gramene out-performed PH207-community method in all categories, and the recall was more than 5-10x higher for GOMAP in both CC and BP categories. GOMAP is the only method that achieved higher or comparable performance to other methods in all three categories.

*F*_*max*_ gives a single number for the comparison of the performance of the three methods. Similar to Recall, no one method showed higher performance in all three GO categories. Gramene showed higher performance in MF and BP categories, but GOMAP had higher *F*_*max*_ in CC category. The higher precision achieved by Gramene edged Gramene ahead of GOMAP in both categories, and higher recall edged GOMAP ahead in the CC category. PH207-community method had lower *F*_*max*_ in all three categories, and especially lower by a significant margin in the CC category. PH07-community method showed comparable although slightly lower performance than GOMAP only in MF category. The performance of the PH207-community method was affected by the lower recall observed in all categories.

### Comparison of to GOMAP the Community and Curated Annotations

A comparison of genes, GO terms, and annotations between the GOMAP dataset and community dataset was performed for B73v4 and PH207 in each GO category. This comparison was restricted to the gold-standard terms to provide biological validity to the data that was being compared. The recall values of less than one observed in all datasets across all GO categories indicate that no method managed to predict all the annotations in the gold-standard dataset (Figure 4). The comparison allowed for the identification of unique genes and GO terms that were only annotated by a particular method. The comparative proportions of the comparisons are presented in Figure 5 and absolute numbers are presented in Table 5.

**Figure 5:**
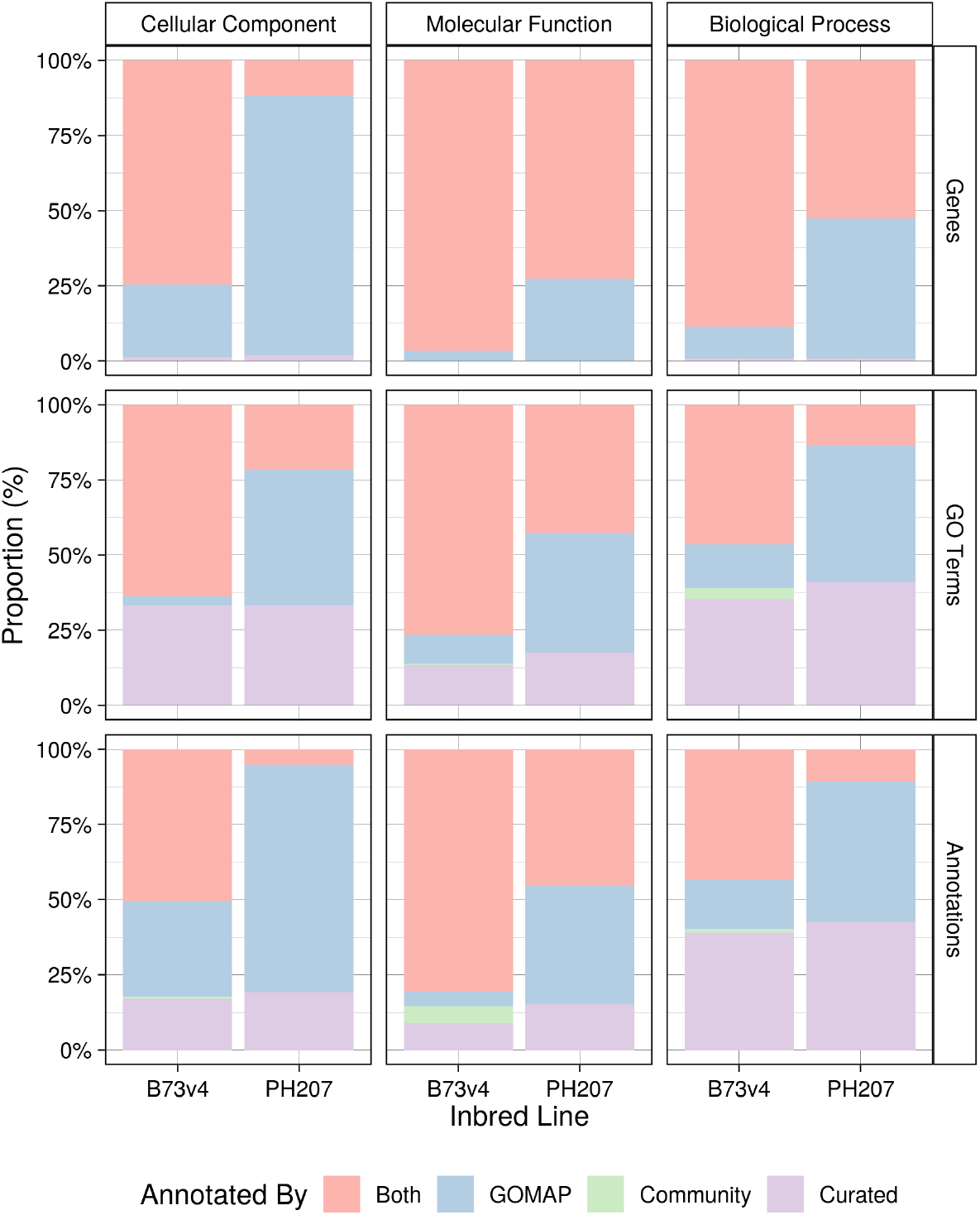
Comparison of the GOMAP and community annotations based on whether gold standard terms were annotated. Left column: Cellular Component. Middle column: Molecular Function. Right column: Biological Process. Top row: Percentage of genes with at least one annotation. Middle row: proportion of unique GO terms recovered. Bottom row: proportion of expanded GO annotations recovered. Gold standard genes or annotations recovered by both the community and GOMAP methods are shown in pink. Those recovered by GOMAP but not the community method are shown in blue. Those recovered by the community annotation but not GOMAP are shown in green. Those annotated in the gold-standard that were not recovered by either method are shown in lavender.

GOMAP has annotated more gold-standard genes in both B73v4 and PH207 across all three GO categories than Gramene and PH207-community methods. The majority of genes have annotation from both GOMAP and Gramene for B73v3, but GOMAP and PH207-community methods have annotated a majority of the genes only in MF and BP categories. Due to higher coverage observed in Gramene and GOMAP, the portion gold-standard genes both is higher in all GO categories compared to the proportion of genes annotated by GOMAP and the PH207-community method. Only a few gold-standard genes from CC and BP categories in B73v4 have been annotated by only Gramene, but a larger number of gold-standard genes in CC and BP have been annotated only by GOMAP. No genes were annotated by the PH207-community method that were not annotated by GOMAP, and a substantially higher proportion of PH207 genes have only been annotated by GOMAP. The same trend is also observed in GO terms annotated by different methods. The majority of GO terms were annotated by both methods for B73v4, but GOMAP annotates more terms to gold-standard genes than Gramene. Gramene has annotated only a few terms in BP and one term in MF that were not annotated by GOMAP to any gold-standard genes. GO terms annotated by the PH207-community method are a subset of GOMAP GO terms, and GOMAP has annotated more than twice the number of GO terms annotated by the PH207-community method to gold-standard genes in both the CC and BP categories. Unfortunately, a proportion of GO terms in the gold-standard data has not been annotated by any method for both inbreds. This number varies among GO categories, but is higher in CC and BP than MF. Next, the comparison was performed using the gold-standard annotations (i.e., Curated Gene-GO term pairs). GOMAP outperforms Gramene in the proportion of gold-standard annotations that are correctly predicted in CC and BP, but Gramene outperforms GOMAP for MF. Although the number of gold-standard annotations in MF that are only predicted by GOMAP (31) are similar to Gramene (35), the *F*_*max*_ difference is significant. The PH207-community annotations are a subset of GOMAP annotations and a substantial number of annotations are only found in GOMAP. This is expected based on the recall values seen in Figure 4. Smaller proportions of gold-standard annotations are not predicted by either method in CC (∼11%) and MF (∼8%) categories, but this number increases to (∼40%) in the BP category.

## Discussion

Over the course of the maize-GAMER project our main goal was to improve the maize GO annotation landscape, and develop a reproducible method for annotating plant genomes. During the GOMAP project, we focused on developing a reproducible and high-throughput pipeline that can produce high-coverage and high-quality plant GO annotations. Furthermore, we also wanted the pipeline to be portable across different systems, and be usable by researchers of different backgrounds with minimal effort. We achieved high-quality and high-coverage annotation by streamlining and generalizing GAMER code. We containerized GOMAP for portability and reproducibility, which decreases the effort needed to run the GOMAP pipeline. Moreover, we parallelized the time-consuming steps and decreased the overall runtime from a few weeks to a few days. Since we released GOMAP, six graduate students have annotated gene functions to thirteen plant genomes over the course of an eight-week rotation, and one undergraduate student annotated the grape genome over the course of a single semester for research course credit [33–47].

The comparison of GOMAP to the community annotations illustrates that the GOMAP datasets are higher coverage than other methods as evidenced by coverage and number of annotations. GOMAP shines in recall due to higher coverage, but this comes at the cost of precision. Moreover, a careful comparison of the gold-standard annotations also confirmed that GOMAP does indeed have a significantly lower number of False Negatives (FNs) than other methods. We can accept the sacrifice in precision as long as the potential False Positives (FPs) fall within an acceptable margin for a high-coverage annotation pipeline such as GOMAP. We have used the current set of maize gold-standard annotations to optimize the balance between precision and recall for GOMAP. The gold-standard data available for maize are incomplete and sparse and inflate the number of FPs. The inflation of FPs leads to underestimation of precision, and under-optimized annotation parameters. At present, the accurate identification of FPs with incomplete gold-standard data has been difficult even for larger-scale efforts such as CAFA. Moreover, the sparse gold-standard data also leads to an inflation of FPs for methods that have higher number of annotations, and in this case the Gramene and GOMAP are affected more than PH207-community approach. PH207-method has higher precision in CC and BP categories, and PH207-community in these two categories have lower number of annotations and coverage. In comparison, both GOMAP and Gramene have lower precision in those categories, indicating that some of the correct predictions have been classified as FPs. We expect the gradual accumulation of gold-standard annotations will not only improve the optimization of annotation methods, but also precision metric calculation.

However, the number of annotations predicted by GOMAP in the BP category is high enough that it is possible that GOMAP is producing more FPs. GOMAP annotations show higher overall recall but that could be at the cost of precision. The BP category is known to be the most difficult to predict based on sequence information alone [7], and this is clearly seen in the performance of GOMAP. For future iterations of GOMAP, improvements to the performance of BP category prediction will be a focus for improvements. The lower specificity values for GOMAP-produced datasets compared to those produced by Gramene are explained by the higher number of GOMAP-only annotations that have lower specificity. This interpretation is suggested by figure S1, which shows more lower-specificity annotations. GOMAP is especially affected with a large number of lower specificity annotations in the BP category. However, when the specificity calculation was restricted to genes annotated by community methods, GOMAP showed higher specificity in CC and comparable specificity for MF (See supplemental figure S2).

Comparison of the methods also indicates that GOMAP annotation quality is comparable to the Gramene method. We designed GOMAP not as a replacement for Gramene but as a supplemental source of annotations. Gramene has been a important resource that provides the plant community with high-quality annotations and invaluable community outreach, and is a federally funded organization. The latest updates from Gramene indicate large-scale curation efforts to improve functional annotations [48]. The curation efforts will improve the annotations of the plant genomes currently available in Gramene, but will not be easily transferable to newly assembled genomes. Unfortunately, Gramene doesn’t include all newly released plant genomes. For example, Three out of the four inbred lines that were annotated in this paper are not currently available in Gramene. We expect GOMAP to allows researchers to annotate their own plant genomes or translated transcriptomes in a high-throughput manner and produce annotations of comparable quality to sophisticated methods employed by Gramene. This reduces the time for functional annotation of newly assembled genomes and leads to better understanding of the sequenced genomes.

The current version of GOMAP focuses on genome-wide functional annotation using multiple methods, some of which are themselves computationally intensive, which results in high computational requirements for the GOMAP system. GOMAP’s component methods including InterProScan and the sequence comparison to the UniProt sequence database significantly contribute to the computational requirements compared to, for example, the simple BLAST searches used by the community to annotate PH207 [13]. It would be interesting to compare computational requirements between GOMAP and Gramene’s annotation pipeline given that both are systems that are reported to use multiple methods. However, the pipeline used by Gramene does not have sufficient documentation to enable anyone outside of outside of Gramene to reproduce their annotations directly. Gramene has evolved over the course of various releases, and incorporates multiple methods such as the Ensembl Compara pipeline for building phylogenies, InterProScan for domain annotation, and cellular localization signals for functional annotation. Dataset content also indicates that Gramene likely inherits GO annotations from Uniprot GOA, InterProScan, curated datasets, and other sources [48]. The lack of detailed documentation precludes a direct computing performance comparison between GOMAP and the Gramene systems, but this is understandable given that the scope of the Gramene project is well beyond GO-based functional annotation for genes [48].

We started developing GOMAP after the first round of the CAFA competition (CAFA1) results had been announced. GAMER and subsequently GOMAP were developed based on three of the top performing CAFA1 methods. Overall performance of CAFA1 methods were better than naive methods such as BLAST or Pfam. We are grateful for the effort to organize the CAFA competitions and the function prediction community for developing these methods for GOMAP. Moreover, the CAFA competition standardized evaluation methods and provided an unbiased and effective method to compare across different annotation methods. While methods that participated in CAFA improved the quality of the predictions, they were not assessed in the context of annotating non-model plants nor for genome-wide performance. GOMAP bridges the gap between the top performing functional annotation methods and adapts them to a plant-specific context. Over the course of the GOMAP project, we also assessed and optimized the quantity of annotations produced for entire genomes. We have seen continuous improvement in the function prediction methods over CAFA2 and CAFA3. The top performing methods of CAFA2 and CAFA3 have improved the quality of the annotations further as evidenced by *Fmax*. We expect GOMAP can be further improved by adding top performing methods from CAFA2 and CAFA3 to the system. Assessing newer tools could also allow us to decouple GOMAP from external methods such as Argot2 and create a self-contained pipeline without sacrificing the quality of annotations produced. As additional features, the next iteration of GOMAP development for customizability and a conda package to improve usability.

## Author’s contributions

K.W. designed the pipeline and the computational implementation and analyzed the data. K.W. and C.J.L-D. wrote the manuscript.

## Acknowledgments

Thanks to: R. Walls and D. Campbell for generating data DOIs and hosting GOMAP data on CyVerse; N. Weeks for helping adapt FANN-GO to use GNU Octave instead of MATLAB; I. Braun, S. Cannon, A. Jain, G. Kandoi, and N. Weeks for testing the GOMAP pipeline and for valuable suggestions. Thanks to K. Chiteri, H. Dostalik, L. Fattel, P. Joshi, H. Vu, D. Psaroudakis, and C. Yanarella for using GOMAP to annotate plant genomes. Finally, thanks to all members of the Dill Plant Informatics and Computational Lab (dill-picl.org) for in-depth discussions about the project and for offering helpful suggestions.

This work has been supported by the XSEDE startup allocation awarded to K.W. and C.J.L-D; funding from the Iowa State University Plant Sciences Institute Faculty Scholars Program to C.J.L-D.; and funding from the National Science Foundation [IOS #1027527] to C.J.L-D.

## Additional Files

Additional file 1 — GOMAP-Supplemental

The supplemental tables are provided as a PDF file that can be opened with the free software.

## Footnotes

[1] http://datacommons.cyverse.org/browse/iplant/home/shared/dillpicl/gomap/GOMAP

[2] https://github.com/Dill-PICL/GOMAP-singularity

[3] https://www.hpc.iastate.edu/guides/condo-2017

[4] https://www.psc.edu/resources/bridges/

[5] https://github.com/wkpalan/GOMAP-maize-analysis

[6] https://maizegdb.org/search/gene/download_gene_xrefs.php?relative=v4

