## Supplemental Tables and Figures for "Gene Ontology Meta Annotator for Plants (GOMAP)"

### GOMAP Supplemental Data

Kokulapalan Wimalanathan

2/23/2021

#### Supplemental Tables

Table ST1: Location of input fasta files for the maize lines

| Inbred Line | Link |
|---|---|
| B73v4 | ftp://ftp.ensemblgenomes.org/pub/plants/release-32/fasta/zea_mays/pep/Zea_mays.AGPv4.pep.all.fa.gz |
| B73v3 | https://ftp.maizegdb.orgGDB/FTP/B73_RefGen_v3/Zea_mays.AGPv3.22.pep.all.fa.gz |
| Mo17 | https://ftp.maizegdb.orgGDB/FTP/Zm-Mo17-REFERENCE-CAU-1.0/Zm00014a.proteins.fa.gz |
| PH207 | https://ftp.maizegdb.orgGDB/FTP/Zm-PH207-REFERENCE_NS-UIUC_UMN-1.0/Zm00008a.protein.fa.gz |
| W22 | https://ftp.maizegdb.orgGDB/FTP/Zm-W22-REFERENCE-NRGENE-2.0/Zm00004b.protein.fa.gz |

Table ST2: Comparison of GOMAP Annotations and Plant-Specific Subset

| | | Cellular Component | | Molecular Function | | Biologial Process | |
|---|---|---|---|---|---|---|---|
| inbred | source | All | Plant-specific | All | Plant-specific | All | Plant-specific |
| B73v3 | GOMAP | 135,251 | 133,272 | 87,953 | 87,267 | 291,855 | 287,199 |
| B73v4 | Community | 35,771 | 35,764 | 49,659 | 49,685 | 44,998 | 45,016 |
| B73v4 | GOMAP | 88,831 | 87,683 | 82,849 | 81,906 | 278,952 | 270,146 |
| Mo17 | GOMAP | 87,573 | 86,482 | 79,755 | 78,892 | 278,043 | 269,274 |
| PH207 | Community | 4,429 | 4,428 | 23,192 | 23,219 | 13,093 | 13,081 |
| PH207 | GOMAP | 90,625 | 89,511 | 86,106 | 85,151 | 288,937 | 278,278 |
| W22 | GOMAP | 95,397 | 94,271 | 85,616 | 84,699 | 290,032 | 280,705 |

Figure S1: Distribution of Specificity for the maize annotations from community and GOMAP

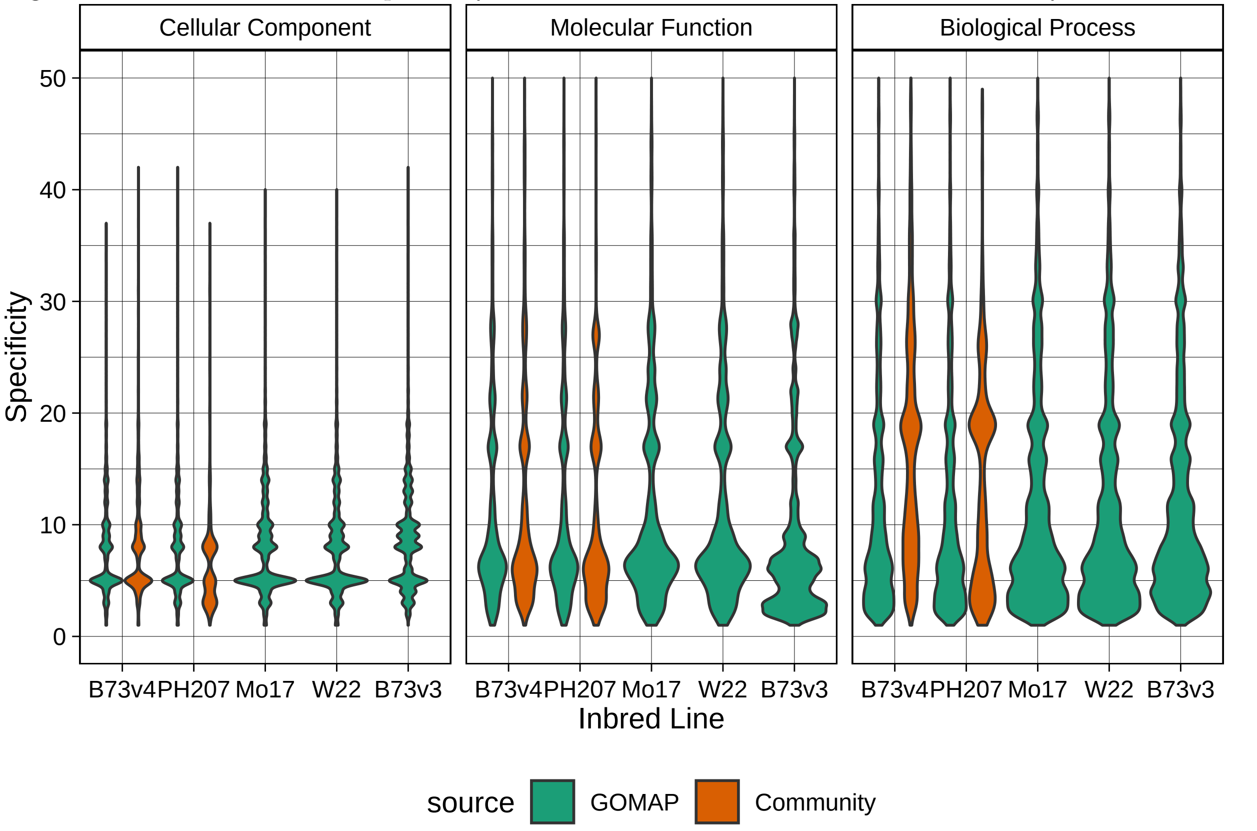


Left column: Cellular Component. Middle column: Molecular Function. Right column: Biological Process. Inbred lines are denoted along the x-axis. The specificity is denoted along y-axis, and the range is limited to less than 50. GOMAP annotations are denoted by green color. Community annotations are denoted by a orange color. The width of the plot along the y-axis indicates the proportion of the annotations that particular specificity value.

Figure S2: Distribution of Specificity for the genes that are shared between community and GOMAP

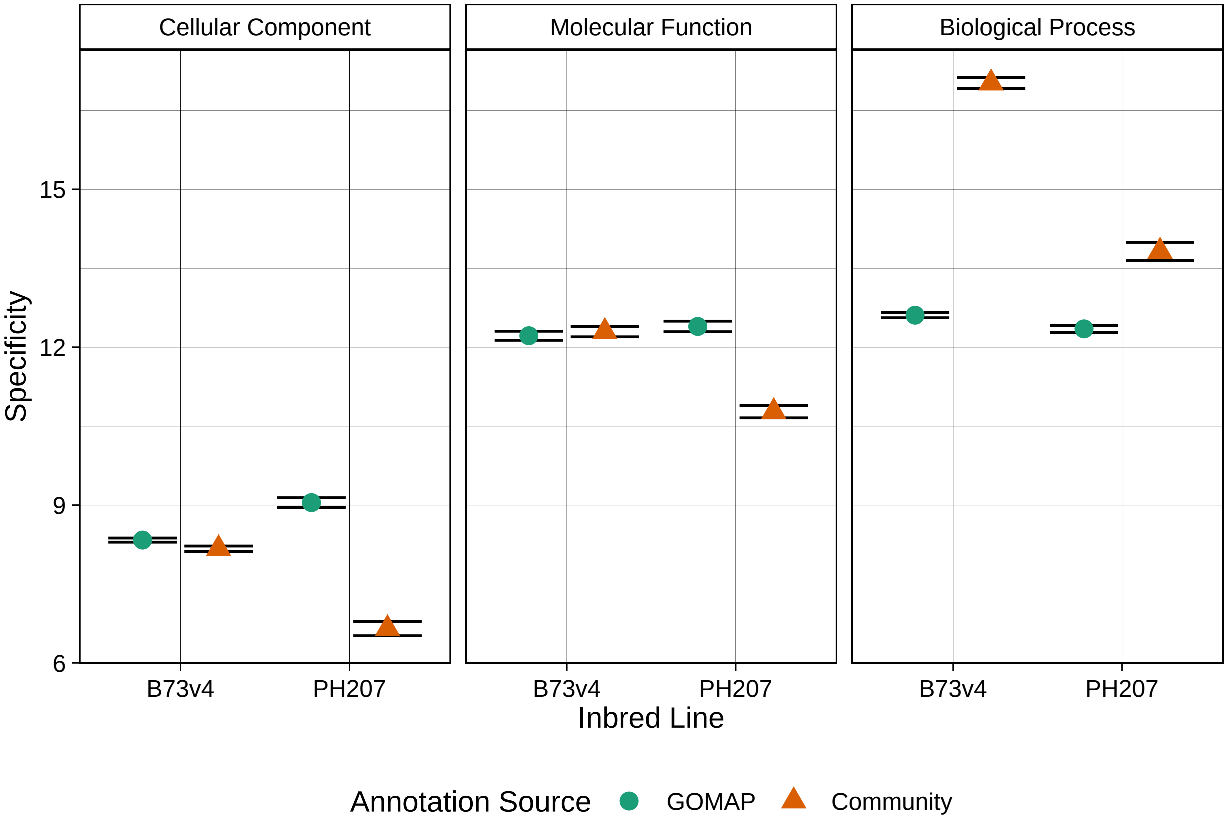


Left column: Cellular Component. Middle column: Molecular Function. Right column: Biological Process. Inbred lines are denoted along the x-axis. The specificity is denoted along y-axis, and the range is limited to less than 50. GOMAP annotations are denoted by green color. Community annotations are denoted by a orange color. The width of the plot along the y-axis indicates the proportion of the annotations that particular specificity value.
